# A genetically encoded single-wavelength sensor for imaging cytosolic and cell surface ATP

**DOI:** 10.1101/385484

**Authors:** Mark Lobas, Jun Nagai, Mira T. Kronschläger, Philip Borden, Jonathan S. Marvin, Loren L. Looger, Baljit S. Khakh

## Abstract

Adenosine 5’ triphosphate (ATP) is a universal intracellular energy source^1^ and an evolutionarily ancient^2^ extracellular signal^3–5^. Here, we report the generation and characterization of single-wavelength genetically encoded fluorescent sensors (iATPSnFRs) for imaging extracellular and cytosolic ATP from insertion of circularly permuted superfolder GFP into the epsilon subunit of F_0_F_1_-ATPase from *Bacillus PS3*. On the cell surface and within the cytosol, iATPSnFR^1.0^ responded to relevant ATP concentrations (30 μM to 3 mM) with fast increases in fluorescence. iATPSnFRs can be genetically targeted to specific cell types and sub-cellular compartments, imaged with standard light microscopes, do not respond to other nucleotides and nucleosides, and when fused with a red fluorescent protein function as ratiometric indicators. iATPSnFRs represent promising new reagents for imaging ATP dynamics.

---

Given widespread roles in energy homeostasis and cell signaling^3–5^, several methods have been used to detect ATP using small-molecule chemical approaches^6^, firefly luciferase^7,8^, ion channel-expressing “sniffer” cells^9–11^, voltammetry^12^, and different types of microelectrodes^13,14^. However, these methods lack spatial resolution, cannot be easily used in tissue slices or *in vivo*, respond to off-target ligands, need unwieldy photon-counting cameras, are imprecise and/or unavoidably damage tissue during probe placement. Importantly, none of these methods can be genetically targeted to specific cells, sub-cellular compartments or whole organisms, and they all lack cellular-scale spatial resolution. Although luciferase is genetically encoded^15^, it requires the exogenous substrate luciferin, addition of which complicates use in tissue slices and *in vivo*. It also saturates at nanomolar ATP, far lower than concentrations expected during ATP signaling (~1 μM to ~1 mM). Additionally, luciferase’s bioluminescent output yields low photon fluxes and rules out cellular-resolution imaging. To address these issues, genetically-encoded fluorescent ATP sensors have been developed, including Perceval and PercevalHR^16,17^, ATeam^18^, and QUEEN^19^. These sensors are valuable, but they have limitations. They either respond to ADP/ATP ratio instead of ATP concentration or are susceptible to optical overlap with cellular sources of auto-fluorescence, and all are incompatible with single-wavelength fluorescence imaging, the workhorse of functional fluorescence microscopy^20^.

Our goals were to: 1) develop an ATP sensor that could be used in routine fluorescence imaging, *e.g.* at GFP’s 488 nm excitation and ~525 nm emission and 2) deploy such a sensor on the cell surface and within cells to image ATP. QUEEN is an excitation ratio sensor, ATeam is a fluorescence resonance energy transfer (FRET) sensor, and the Perceval sensors are fluorescence lifetime indicators – all of these methods require customized equipment, substantially slow down imaging rates, complicate use with other reagents, and typically have lower signal-to-noise ratio (SNR) than single fluorescence-wavelength intensity indicators. After unsuccessful attempts to modify ATP P2X receptors^21^ and the bacterial regulatory protein GlnK1 (as in Perceval^16^) to suit our needs, we turned to microbial F0F1-ATP synthase epsilon subunits^22^, which have been adapted to create the FRET sensor ATeam^18^ and the excitation ratio sensor QUEEN^18,19^. The epsilon subunit is predicted to undergo a large conformational change^22^ upon binding ATP (Fig. 1a), and the homologue from the thermophilic bacterium *Bacillus PS3* displays appropriate ATP-binding sensitivity^23^. The epsilon subunit comprises eight N-terminal β-strands followed by two C-terminal α-helices (Fig. 1a), which extend away from the β-strands in the absence of ATP, but upon binding cradle ATP up against the β-strands.

**Figure 1:**
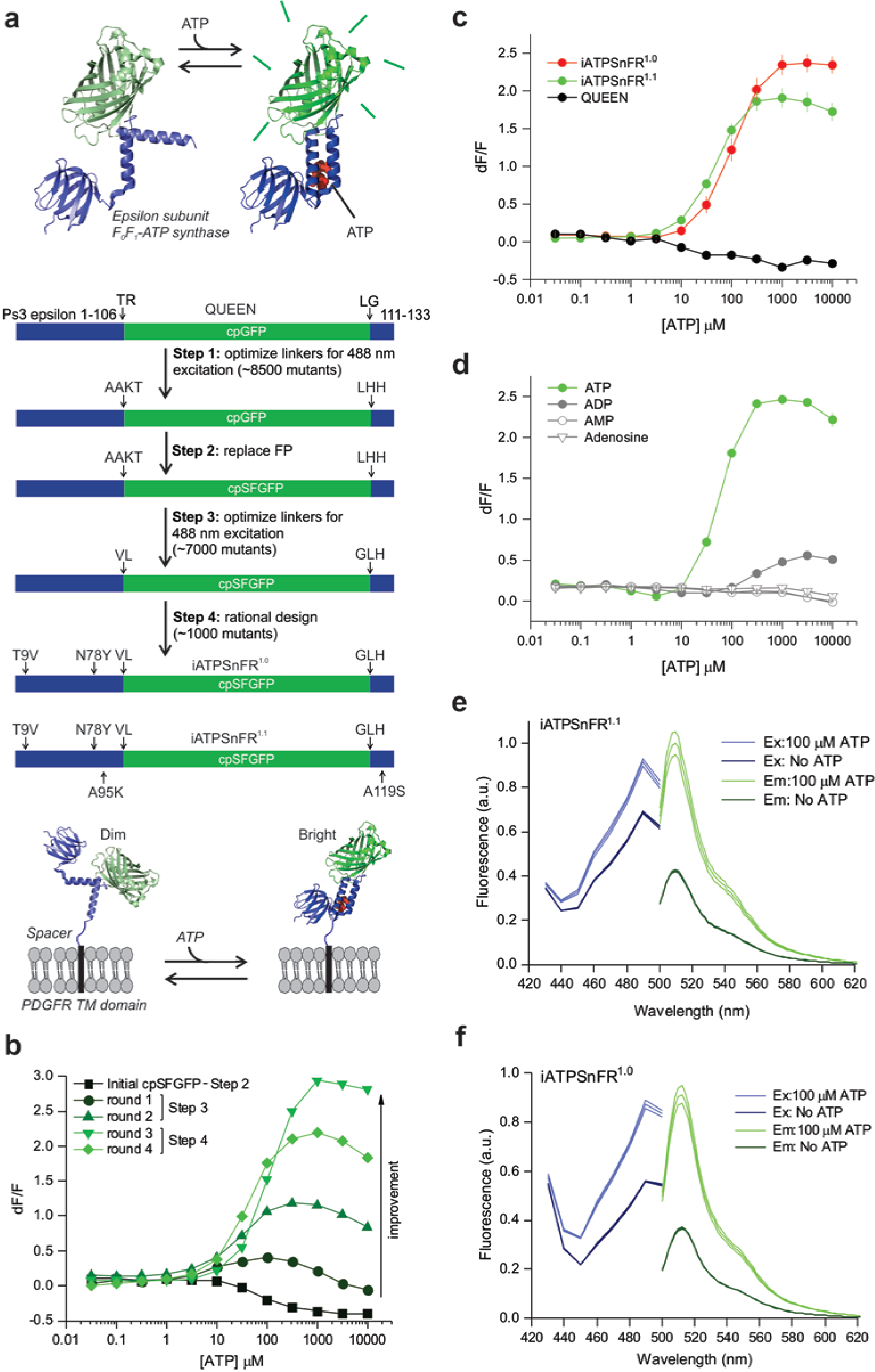
Design and optimization of a single-wavelength ATP sensor. (**a**) Schematic showing the design and workflow used to optimize QUEEN-7µ into a single-wavelength ATP sensor with the goal of displaying the sensor on the surface of cells. (**b**) Dose-response curves of iATPSnFR over several successive rounds of mutagenesis (ex: 488 nm, em: 515 nm). Fluorescence quenching at very high ATP concentrations can be observed in addition to binding-dependent increases. (**c**) Dose-response curves for purified QUEEN-7µ, iATPSnFR^1.0^ and iATPSnFR^1.1^. (**d**) Dose-response curves of purified iATPSnFR^1.1^ to ATP, ADP, AMP, and adenosine. (**e-f**) Excitation and emission spectra for iATPSnFR^1.0^ and iATPSnFR^1.1^. In some cases, the error bars for the s.e.m. are smaller than the symbols used for the mean.

Circularly permuted (cpGFP)^24^ was inserted between the two α-helices of the epsilon subunit after residue 107 with the expectation that the epsilon subunit conformational change might alter fluorescence. The first linker (L1) initially comprised Thr-Arg, with the second linker (L2) Leu-Gly (Fig. 1a). Based on our past experience with the glutamate sensor iGluSnFR^25,26^, we began mutating residues in the linkers (~8500 colonies screened) to develop sensors with large ATP-dependent fluorescence intensity increases (dF/F). The best variant from this screen had a large change in fluorescence (maximum dF/F of ~3.9). However, it failed to express on the surface of HEK293 cells when cloned into the pDisplay mammalian expression vector, which uses an IgG secretion signal and a platelet-derived growth factor receptor (PDGFR) transmembrane domain to anchor it to the membrane. We reasoned that a more stable form of GFP might improve folding and trafficking, and thus cloned circularly permuted superfolder GFP^24^ (cpSFGFP) in place of cpGFP. Replacing cpGFP with cpSFGFP remedied the surface trafficking in HEK293 cells (see later section), but greatly diminished ATP-evoked changes in fluorescence. To correct this, we re-optimized L1 and L2 for the cpSFGFP construct by mutating amino acids in the linkers (and slightly changing their length; ~7000 colonies screened; Fig. 1a & b). We also mutated amino acids (Thr9Val and Asn78Tyr) predicted from molecular modeling to decrease dimer formation. Through this process, we developed two sensors that display large ATP-dependent increases in fluorescence (Fig. 1a & b). In the sensor we termed iATPSnFR^1.0^, the L1 linker was changed from Thr-Arg to Val-Leu, and L2 from Leu-Gly to Gly-Leu-His. We developed a second sensor (iATPSnFR^1.1^) with improved sensitivity by mutating amino acids near the ATP-binding pocket. iATPSnFR^1.1^ differs from iATPSnFR^1.0^ by two mutations (Ala95Lys and Ala119Ser; Fig. 1a; **Supp. Fig. 1**). Both iATPSnFR^1.0^ and iATPSnFR^1.1^ show marked improvement over QUEEN-7µ, which does not function as a single-wavelength sensor (Fig. 1c). Furthermore, inserting cpSFGFP into Queen did not result in a sensor with ATP-evoked fluorescence increases. Purified iATPSnFR^1.0^ had a maximum dF/F of ~2.4 and an EC_50_ of ~120 µM, whereas purified iATPSnFR^1.1^ had a maximum dF/F of ~1.9 and an EC_50_ of ~50 µM (Fig 1c). Purified iATPSnFRs were not sensitive to ADP, AMP, or adenosine at concentrations equivalent to ATP (Fig. 1d). Both proteins displayed similar fluorescence spectra (peak excitation 490 nm, peak emission 512 nm; Fig. 1e & f). In the presence of ATP, a large increase in peak excitation and emission was observed. Thus, our bacterial-based screen yielded a genetically encoded ATP sensor with micromolar sensitivity, large 488 nm-excited emission intensity dF/F, and little sensitivity to ATP metabolites (Fig. 1).

In HEK293 cells co-transfected with cytoplasmic mCherry and membrane-displayed iATPSnFR^1.0^ or iATPSnFR^1.1^ (Fig. 1a), there was clear iATPSnFR fluorescence at the cell edges. In contrast, fluorescence was mostly absent from the cytoplasm, indicating proper membrane trafficking (Fig. 2a & 2c). Furthermore, application of 1 mM ATP resulted in increased fluorescence of iATPSnFR^1.0^ and iATPSnFR^1.1^ at the cell edges (Fig. 2b & 2d). Both iATPSnFR^1.0^ and iATPSnFR^1.1^ displayed concentration-dependent increases in fluorescence that were rapid, maintained in ATP, and that returned to baseline upon ATP washout (Fig. 2e; kinetics below). iATPSnFR^1.0^ displayed an EC_50_ of ~350 µM with a maximum dF/F of ~1.0 (Fig. 2f). Similarly, iATPSnFR^1.1^ displayed an EC_50_ of ~140 µM with a maximum dF/F of ~0.9 (Fig. 2g). Thus, there was a ~3-fold loss of ATP sensitivity and decrease in peak response in both sensors when they were extracellularly displayed compared to the soluble proteins (similar effects are seen with other membrane-displayed sensors^25–27^). During these experiments, we noticed that cells expressing iATPSnFR^1.1^ were dimmer than those expressing iATPSnFR^1.0^, and we found lower expression levels of iATPSnFR^1.1^ in relation to iATPSnFR^1.0^ (Fig. 2h). Thus, iATPSnFR^1.1^ has a lower ATP EC_50_ (more sensitive), but is not as well expressed as iATPSnFR^1.0^. iATPSnFR^1.1^ also showed slightly more intracellular accumulation than iATPSnFR^1.0^ (Fig. 1c).

**Figure 2:**
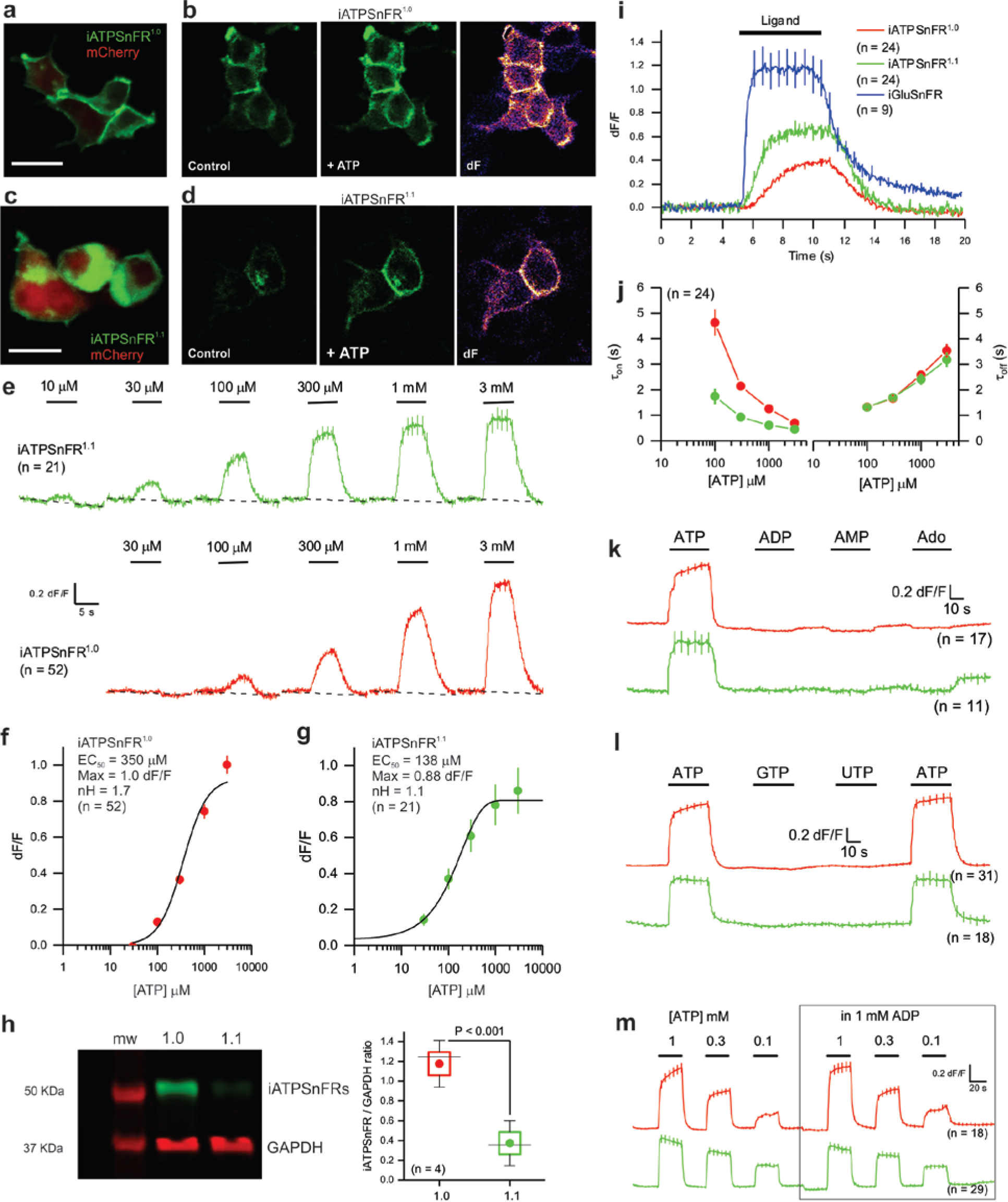
Characterization of cell surface iATPSnFRs in HEK293 cells. (**a**) Single planes from a confocal z-stack of transiently transfected HEK293 cells expressing cytoplasmic mCherry and membrane-displayed iATPSnFR^1.0^. (**b**) Confocal images of transiently transfected HEK293 cells expressing iATPSnFR^1.0^ before ATP application (control) and during a 1 mM ATP application (+ATP), as well as the change in fluorescence (dF). (**c-d**) As in a-b, but for iATPSnFR^1.1^ transiently expressed in HEK293 cells. (**e**) Average traces for iATPSnFR^1.1^ (green) and iATPSnFR^1.0^ (red) over a range of concentrations from 10 µM to 3 mM. (**f & g**) Dose-response curves for iATPSnFR^1.0^ and iATPSnFR^1.1^ displayed on the surface of HEK293 cells. Sensitivity (EC_50_), total dynamic range (Max) and cooperativity (nH) shown. (**h**) Representative Western blot from HEK293 cells transfected with iATPSnFR^1.0^ or iATPSnFR^1.1^ probed for GFP (iATPSnFRs) and GAPDH (loading control). The graph shows the quantification of iATPSnFR expression from four Western blots normalized to GAPDH loading control. The circle represents the mean, the box the s.e.m, the whiskers the s.d and the horizontal line the median. (**i**) Traces for cell-displayed iATPSnFR^1.0^, iATPSnFR^1.1^ and iGluSnFR kinetics from fast-solution change experiments. We could change solutions in under ~10 ms. (**j**) Shows the on and off for iATPSnFR^1.0^ and iATPSnFR^1.1^ at various ATP concentrations. (**k**) Shows the response of iATPSnFR^1.0^ and iATPSnFR^1.1^ to ATP, ADP, AMP, and adenosine (1 mM applications). (**l**) The response of iATPSnFR^1.0^ and iATPSnFR^1.1^ to ATP, GTP, and UTP (1 mM applications). (**m**) Traces of iATPSnFR^1.0^ and iATPSnFR^1.1^ with three different concentrations of ATP (1 mM, 0.3 mM, and 0.1 mM) with and without 1 mM ADP. ADP has little effect on iATPSnFR responses. Scale bars are 30 µm. In some cases, the error bars for the s.e.m. are smaller than the symbols used to represent the mean.

Using a fast solution switcher, we assessed the response kinetics of cell-displayed iATPSnFR^1.0^ and iATPSnFR^1.1^ in relation to iGluSnFR, which displays very fast on-and off-rates25,28. The time constants for fluorescence increases (τ_on_) were dose-dependent and significantly faster for iATPSnFR^1.1^ than iATPSnFR^1.0^ (Fig. 2i & j). In contrast, the time constants for the return to baseline (τ_off_) of the ATP-dependent fluorescence increase were not different between the two constructs, implying that the higher equilibrium EC_50_ of iATPSnFR^1.0^ was due to slower association of ATP with the sensor or due to slower subsequent coupling of ATP binding to fluorescence change. In comparison, iGluSnFR was significantly faster than both iATPSnFRs (Fig. 2i & j).

ATP is sequentially degraded to ADP, AMP, and adenosine by ATPases and other ATP-utilizing enzymes in the cytosol and on the cell surface. In accordance with the bacterial lysate data (Fig. 1d), cell surface iATPSnFRs were not sensitive to ADP, AMP, or adenosine at 1 mM (Fig. 2k). Furthermore, iATPSnFRs were not sensitive to 1 mM GTP or 1 mM UTP (Fig. 2l), other nucleoside triphosphates found within the cytosol and in the extracellular space. Since the extracellular space is expected to contain a mixture of ATP and ADP, we evaluated if 1 mM ADP affected the response to 0.1, 0.3 and 1 mM ATP for iATPSnFR^1.0^ and iATPSnFR^1.1^. It did not (Fig. 2m). Thus, iATPSnFRs sense ATP with negligible responses to degradative products or to common competing nucleotides and nucleosides.

For characterization in astrocytes and neurons we focused on iATPSnFR^1.0^, given its better expression. We co-expressed iATPSnFR^1.0^ with mCherry in cultured U373MG astroglia and observed good cell-surface trafficking (Fig. 3a), although there was also an intracellular pool of the sensor. We confirmed that iATPSnFR^1.0^ responded to 300 µM ATP applications with fluorescence increases (Fig. 3b). Similar data were obtained by expressing iATPSnFR^1.0^ in primary cultures of rat hippocampal neurons (Fig. 3c). For U373MG astroglia, we determined the iATPSnFR^1.0^ EC_50_ to be 400 µM with a maximum dF/F of 1.1 (Fig. 3d), similar to the HEK293 cell data. In hippocampal neurons, iATPSnFR^1.0^ properly localized to membrane processes, and application of ~1 mM ATP evoked increases in fluorescence. In hippocampal neurons, the ATP EC_50_ was 630 µM and the maximum dF/F was ~1.5 (Fig. 3e). Hence, our experiences with iATPSnFR^1.0^ in solution, in HEK293 cells, in astroglia and in hippocampal neurons demonstrate that the weakening of ATP binding and the decrease in dF/F resulting from tethering the sensor to the outside of the membrane bilayer is independent of the host cell. We have observed altered sensitivity with other sensors on the membrane^25–27^; we speculate that it results from steric restriction from lipid head groups and membrane proteins.

**Figure 3:**
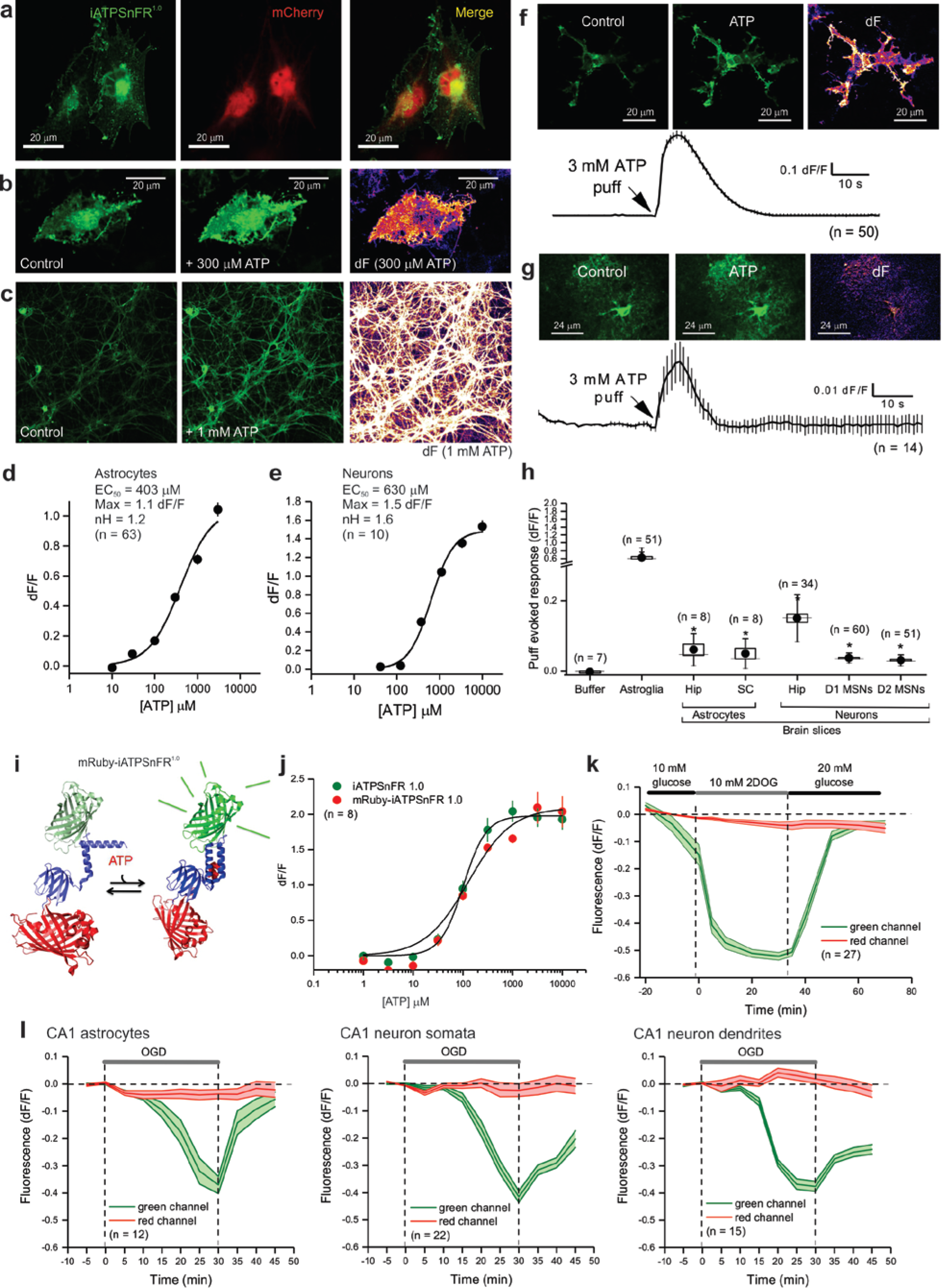
Characterization of cell surface and cytosolic iATPSnFR^1.0^ in astrocytes and neurons. (**a**) A single plane from a confocal z-stack of a U373MG cell expressing cytoplasmic mCherry and membrane-displayed iATPSnFR^1.0^. (**b**) Confocal images of U373MG cells expressing iATPSnFR^1.0^ before ATP application (control) and during a 300 µM ATP application (+ATP), as well as the change in fluorescence (dF). (**c**) Confocal images of hippocampal neurons before ATP application (control) and during 1 mM ATP application, as well as the change in fluorescence (dF). (**d & e**) Dose-response curves for iATPSnFR^1.0^ when displayed on the surface of U373MG astroglia and hippocampal neurons. (**f**) A confocal image of a U373MG astroglia cell transfected with iATPSnFR^1.0^ before a 3 mM puff of ATP (control), during the puff (ATP), as well as the change in fluorescence (dF). An average trace from all cells with the s.e.m. is also shown. (**g**) Representative hippocampal astrocyte in acute brain slice expressing iATPSnFR^1.0^ before the application of ATP (control), during the application of 3 mM ATP (ATP), and the dF. An average trace from all cells with the s.e.m. is also shown. (**h**) Bar graph summary of the effect of 3 mM ATP on iATPSnFR^1.0^ expressed in the cells indicated. The circle represents the mean, the box the s.e.m, the whiskers the s.d. and the horizontal line the median. The asterix indicates the data that were statistically different to the buffer puff. Abbreviations: hip, hippocampus; SC, spinal cord; MSNs, medium spiny neurons. Note broken y-axis. **(i)** Cartoon schematic of mRuby-iATPSnFR^1.0^: ATP binding is schematized to show an increase in cpSFGFP fluorescence, whereas mRuby fluorescence remains unchanged. **(j)** Dose-response curves for iATPSnFR^1.0^ in relation to mRuby-iATPSnFR^1.0^ in HEK293 cell lysates. **(k)** Changes in fluorescence of mRuby-iATPSnFR^1.0^ (green and red channels) before, during and after inhibition of glycolysis with 2DOG in HEK293 cells. **(l)** Changes in fluorescence of mRuby-iATPSnFR^1.0^ (green and red channels) during oxygen glucose deprivation in hippocampal brain slices. Data from astrocytes as well as CA1 pyramidal neuronal cell bodies and dendrites are shown. In some cases, the error bars for the s.e.m. are smaller than the symbols used to represent the mean. N numbers are stated in the figure panels.

We generated adeno-associated virus (AAV2/5) to express iATPSnFR^1.0^ specifically in astrocytes using the *GfaABC_1_D* promoter^29^. Following *in vivo* microinjections into the hippocampus, iATPSnFR^1.0^ co-localized with S100β-positive astrocytes, but not NeuN-positive neurons (**Supp. Fig. 2**). Bath application of ATP to multicellular preparations such as brain slices is fraught with problems caused by the rapid hydrolysis of ATP by cell surface ectoATPases. In order to minimize such effects, we puffed ATP *via* a glass pipette with a Picospritzer. Puffing 3 mM ATP (for 5 s) onto U373MG astroglia resulted in a ~0.7 dF/F (Fig. 3f). Similar, albeit smaller, responses were observed when we applied ATP to astrocytes expressing iATPSnFR^1.0^ in hippocampus and spinal cord slices or to D1 or D2 medium spiny neurons expressing iATPSnFR in striatum (Fig. 3g, h). Taken together, these data show that iATPSnFR can be expressed and responds reliably on the surface of cells following *in vivo* expression.

We next explored the use of cytosolic iATPSnFR^1.0^ to follow intracellular ATP, which ranges from 1-3 mM in mammalian cells^30^. We expressed cytosolic iATPSnFR^1.0^ in U373MG astroglia and detected a significant drop in intracellular fluorescence upon switching to 0 mM glucose (**Supp. Fig. 3a**), or by applying 10 mM 2-deoxyglucose (2DOG; **Supp. Fig. 3b**) – an inhibitor of glycolysis^31^. Similarly, following *in vivo* expression with AAVs, we measured a significant drop in intracellular ATP levels as reported by the fluorescence of iATPSnFR^1.0^ in hippocampal astrocytes in 0 mM glucose (**Supp. Fig. 3c**; the fluorescence of co-expressed tdTomato did not change, arguing against changes in cell shape and volume, or other non-specific artifacts, as the cause). We repeated these experiments for hippocampal astrocytes and CA1 pyramidal neurons, both at somata and along processes (**Supp. Fig. 3d**): oxygen-glucose deprivation (OGD) caused a substantial drop in intracellular ATP levels. We found that fluorescence decreases of astrocytes were twice that of neurons (**Supp. Fig 3d**). Such responses were completely reversible for astrocytes, but only partly reversible for neurons (**Supp. Fig. 3d**). Furthermore, it took longer for decreases in astrocyte ATP to begin (~10 min in OGD). However, once ATP began to decrease in astrocytes, it took 8.0 ± 0.5 min (n = 37) to decrease by 50%, whereas in CA1 pyramidal neuron somata and dendrites it took 13.5 ± 0.8 min (n = 26) and 17.4 ± 0.9 min (n = 27), respectively (*P* < 0.05 by ANOVA and Dunn *post hoc* tests). Based on the titrations in Fig. 2, we estimate that ATP levels fell by ~0.5 mM from > 1mM. Thus during glucose deprivation, ATP levels in astrocytes fall quickly, but recover completely upon restoration of glucose. In contrast, neuronal ATP levels fall slowly and fail to recover to normal levels under the identical conditions.

We fused the red fluorescent protein mRuby^32^ to the N-terminus of iATPSnFR^1.0^ to generate a ratiometric sensor (Fig. 3i; mRuby-iATPSnFR^1.0^). The mRuby-iATPSnFR^1.0^ construct was identical to iATPSnFR^1.0^ in terms of its sensitivity and ATP-evoked dF/F (Fig. 3j). In HEK293 cells, mRuby-iATPSnFR^1.0^ displayed clear changes in green fluorescence in the presence of 2DOG, whereas the red fluorescence of mRuby did not (Fig. 3k; **Supp. Fig. 4**). Similarly, following *in vivo* expression with AAVs, we measured a significant drop in intracellular ATP as reported by the fluorescence of iATPSnFR^1.0^ in hippocampal astrocytes and pyramidal neuron somata and dendrites upon switching to solutions that mimic ischemia (*i.e.* OGD), whereas there was no change in mRuby (Fig. 3l; **Supp. Fig. 5**). Interestingly, ATP levels returned back to normal levels upon reapplication of physiological buffers in astrocytes, but remained low in neuronal somata and dendrites (Fig. 3l). Hence, mRuby-iATPSnFR^1.0^ can be used as a ratiometric sensor of cell specific intracellular ATP levels.

As expected for GFP-based sensors, the fluorescence of both iATPSnFR^1.0^ and mRuby-iATPSnFR^1.0^ was sensitive to pH, with dF/F decreasing from ~2 at pH 8.0 to 0.6 at pH 6.0, but affinity for ATP remained unchanged by pH (**Supp. Fig. 6a,b**). iATPSnFR^1.0^ is not fluorescent at pH 4.0. Between pH 6.0 and 9.0, fluorescence of ligand-free iATPSnFR^1.0^ was relatively stable, but strongly attenuated by acidic pH in the ATP-bound state (pK_a_ of ~6.5; **Supp. Fig. 6c**). The kinetics for ATP binding to iATPSnFR^1.0^ in solution did not follow typical pseudo-first order behavior (**Supp. Fig 6d**), but were instead consistent with a mechanism in which both higher and lower fluorescent states occur (potentially resulting from monomeric and dimeric states).

We used the bright, stable cpSFGFP scaffold, rational design and library screening to direct the selection of ATP sensors with desired characteristics. We characterised the ATP sensors to monitor cell surface and cytosolic ATP levels with excellent specificity. The current sensors may be useful to study cytosolic ATP levels and signals associated with P2X7 receptors, which respond to ATP in the hundreds of micromolar to millimolar range^33^. They may also be useful for diagnostics to sense high ATP levels in tumors^34^. To better detect small ATP fluctuations in living animals, the sensitivity, dF/F and kinetics should be improved. iATPSnFRs portend the direct imaging of purinergic signaling^5^ throughout the body of transgenic animals. Our constructs are available (**Supp. Table 1**) to allow such studies to commence.

## Acknowledgements

Supported by the NIH (MH104069 and NS100525 to BSK) and Howard Hughes Medical Institute, Janelia Research Campus (to LLL). Looger and Khakh collaborations were part of the Janelia Visiting Scientist Program. JN was supported by a JSPS Overseas Research Fellowship (H28-729). Thanks to Gary Yellen (Harvard) for discussions. Thanks to Joshua Burda for performing spinal cord AAV microinjections and to Michael V. Sofroniew for sharing equipment.

## Author contributions

ML performed molecular biology and imaging: JN, MTK, PB, and JSM helped. JSM and LLL supervised sensor design and optimization. BSK and LLL conceived the project. BSK helped analyze data, assembled figures and wrote the paper: everyone contributed.

## Materials and methods

provided in Supplementary Information

